# Oral Microbiota Interactions with Titanium Implants: A pilot in-vivo and in-vitro study on the impact of Peri-Implantitis

**DOI:** 10.1101/2025.03.09.642253

**Authors:** Priyadharshini Sekar, Zuha Rizvi, Nizam Abdullah, A R Samsudin, Waad Kheder

## Abstract

**Introduction:** Dental implant therapy is widely recognized as the best solution for replacing missing teeth. However, it still has a 1.9-11% failure rate. Oral microbiota is suspected to play a significant role in implant failure and peri-implant infections. Hence the impact of oral microbiota on titanium dental implants, particularly in the context of dysbiosis and peri-implant diseases, was investigated by in-vivo and in-vitro methods in this study.

**Materials and Methods:** This pilot study aimed to investigate the role of oral microbiota in peri-implant diseases associated with titanium implants. For the in-vivo study, oral microbiota was collected from titanium and hydroxyapatite (tooth-mimicking) discs placed in a custom-made intra-oral device worn by four healthy subjects. Biofilm formation, pathobionts, and bacterial diversity were assessed using DAPI staining, qPCR, and next generation sequencing (NGS)-16S rRNA sequencing. For the in-vitro study scanning electron microscopy was employed to examine the effect of oral pathogens on titanium implants.

**Results:** The study found enhanced biofilm formation on titanium implants compared to controls (p < 0.0002). Systematic colonization by *Streptococcus oralis*, *Actinomyces naeslundii*, *Veillonella parvula*, *Fusobacterium nucleatum*, *Porphyromonas gingivalis*, and *Aggregatibacter actinomycetemcomitans* was observed. An abundance of Firmicutes and Bacteroidetes, with a decrease in Proteobacteria on titanium implants was observed by NGS. SEM showed corrosion and damage to titanium implants caused by oral pathogens.

**Conclusion:** The results demonstrate that the use of titanium-based dental implants promotes oral microbiota dysbiosis, tipping the scale towards oral pathogens, which in turn contributes to the damage of the titanium implant surface. Increased biofilm formation of periodontal pathogens and microbial dysbiosis may play a role in implant failure and peri-implant diseases.

## Introduction

Titanium-based dental implants (TDI) have effectively restored missing teeth and supported oral prostheses for over five decades due to their biocompatibility, corrosion resistance, and mechanical strength. However, clinical failures like mucositis and peri-implantitis can occur due to poor oral hygiene, overloaded prostheses, leaching of titanium ions, and infections by oral commensals [1]. The pathogenesis of peri-implantitis remains under investigation [2], but implant failure often starts with titanium surface exposure, leading to microbial attachment and biofilm formation resistant to standard oral care.

The physicochemical properties and topology of the implant surface, along with the characteristics of oral microorganisms and the duration of exposure to the oral environment, influence biofilm establishment [3]. Bacterial adhesion to dental materials is the primary cause of infections and understanding the relationship between these factors and biofilm formation may help to understand the causes of dental implant failure [4]. Additionally, bacterial metabolites can stimulate inflammation and damage the implant surface, leading to corrosion and oxidative stress, which further promote a proinflammatory immune response [5,6]. Investigating the interactions between titanium dental implant surfaces and oral microbiota is essential.

Numerous in-vitro studies on microbial adherence and colonization of biomaterial surfaces show varying results compared to microbiota isolated from failed dental implants in the oral cavity [7,8,9]. Factors such as prophylactic antibiotic use, physiological disturbances, microbiological sampling methods, and oral hygiene may influence the microbiota composition. A controlled in-vivo method using pre-fabricated intraoral devices with test material substrates allows reliable and reproducible biofilm collection and analysis [10,11] This study aimed to investigate oral microbiota dynamics and interactions on TDI surfaces in both in-vitro and in-vivo models.

## Materials and Methods

### In vivo studies on the impact of TDI on oral microbiota

#### Study participants

Four healthy volunteers (ages 19-65) were recruited from the University Dental Hospital and RIMHS, Sharjah, for this quasi-experimental pilot study, following informed written consent and approval from the institute research ethics committee. Participants had good oral hygiene and were excluded if they were pregnant, smokers, had any systemic illness, active periodontal disease, carious lesions, or had used antibiotics, mouthwashes, or other medications in the past six months.

#### Design and use of a custom-fit intra-oral device for harvesting oral microbiota

Volunteers had alginate impressions of their maxillary dental arch made using stock trays, which were then used to create customized acrylic intraoral devices [12]. Each device featured six drilled pits (5 mm diameter) on the buccal side. Six sterilized plastic wells from a microtiter plate were secured in the pits, with each well containing either hydroxyapatite (HA) pellets or segments of titanium implants. This setup included an empty well (control for gum microbial sampling), a well with a titanium disc (3×3 mm), and a well with a HA pellet (2×2 mm), resembling tooth enamel (Fig. 1). Participants wore the intraoral device for 6 consecutive hours to allow oral microbiota colonization on the substrate inside the wells. After this period, the devices were removed under aseptic conditions in the laboratory, and the oral microbiota was collected and stored in phosphate buffered saline (PBS) at 4°C for further processing.

**Fig. 1:**
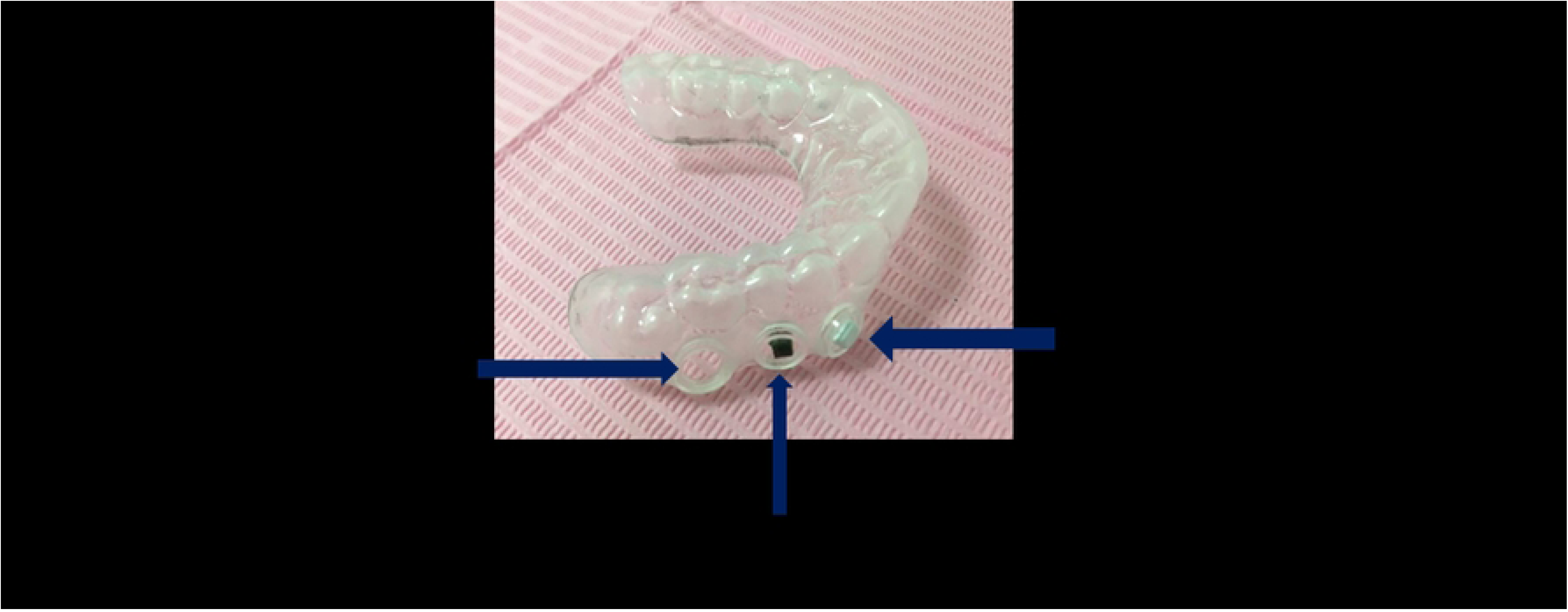
The dental device fitted with three holes on either side, fitted with plastic wells.

#### Microscopical examination of biofilms by DAPI staining and Fluorescence microscopy

TDI, HA pellet, and empty well were assessed for microbial biofilm formation using DAPI (4’,6-diamidino-2-phenylindole) staining (ab104139, Abcam). Each sample was covered with 1 mL of DAPI solution (1 mg/mL) for 10 minutes in darkness, then rinsed with distilled water. Bacterial biofilms were visualized by fluorescence microscopy. Bacterial counts were quantified at 1000x magnification (Axioskop II, Zeiss, Germany) using a DAPI filter set. The number of cells in 10 randomized ocular grid fields per sample was counted, with the ocular grid area (0.0156 mm²) used to estimate cells per cm².

#### DNA extraction

The TDI were rinsed twice in 2 mL of sterile PBS (10 seconds per rinse) to remove non-adherent bacteria. These were then sonicated for 10 minutes at 37°C and 50 Hz to detach bacterial cells. DNA was extracted using the QIAamp DNA microbiome kit (Cat. No. 51704, Qiagen), quantified, and assessed for purity with a NanoDrop2000 (Thermo Scientific, USA) before being stored at −20°C. This DNA was used for both Real-Time quantitative PCR(qPCR) and Next Generation Sequencing (NGS)

#### Real-Time PCR targeting 16SrRNA for identification of bacterial colonizers and statistical analysis

*Streptococcus oralis, Actinomyces naeslundii*, *Veillonella parvula*, *Fusobacterium nucleatum Porphyromonas gingivalis* and *Aggregatibacter actinomycetemcomitans* were detected by Real-Time qPCR using primers as described by Sánchez et al [13]. The qPCR was conducted in a 20 µL reaction mixture, containing optimal primer concentrations, along with 5 µL of DNA from samples. The 16S rRNA gene served as the internal control, and 5 µL of sterile water was used as a negative control (no template control, NTC). The fold expression of each bacterial species was calculated by the double delta Ct method and expressed in log10 values, with statistical graphs generated using GraphPad Prism (version 5, Boston, Massachusetts, USA, www.graphpad.com). Additionally, using the same software, statistical analysis was performed by paired t-test.

#### NGS for studying the difference in bacterial diversity between the TDI and HA

For 16S rRNA NGS sequencing, 100 ng to 1 µg of genomic DNA in elution buffer (minimum concentration of 5 ng/µL) was required. A 50 ng DNA sample was used to amplify the variable regions of 16S rRNA with Phusion polymerase for 25-35 PCR cycles. An amplified library was prepared for Illumina sequencing using primers (319F/806R) targeting the V3 and V4 regions of 16S rRNA (approximately 469 bp). After one PCR cycle, sequencing adapters and barcodes were added for further amplification. The Illumina cBot system generated clusters for sequencing on the MiSeq platform. The raw sequence data were analyzed with QIIME Version 1.7.0-dev to assess alpha diversity among the samples.

### In vitro studies on the effect of oral microbiota on titanium dental implants

An in-vitro experiment was conducted to study the colonization dynamics of early and late oral bacteria on titanium-based dental implants. Sterile titanium implants were submerged in Brain heart infusion broth (BHIB) with two set of combinations of early and late bacterial colonizers. The implants were incubated at 37°C for 3 months, with regular medium changes.

- **Set 1:** Lactobacillus sp. and *Streptococcus sp.* (initial colonizers)
- **Set 2:** Streptococcus mutans (initial colonizer), *Porphyromonas gingivalis*, and *Aggregatibacter actinomycetemcomitans* (late colonizers)
- **Control:** A separate set of sterile titanium implants was submerged in BHIB and incubated at 37°C without any bacteria for three months.

After the incubation period, scanning electron microscopy (SEM) imaging was performed to examine the surface characteristics of the titanium implants.

## Results

### In vivo studies on the impact of TDI by oral microbiota

#### Screening of bacterial biofilm by fluorescent microscopy with DAPI staining

DAPI, a fluorescent stain targeting cellular nuclei, was utilized to visualize in vivo biofilms formed on the three distinct materials. Enumeration was feasible in the empty well (representing gum control) (Fig-2a). The bacterial biofilm on HA, which represents the tooth surface, displayed notable density but still enumeration of bacteria was possible (Fig-2b). In contrast, the stained bacterial biofilm appeared dense and thick on the TDI, rendering enumeration impractical under fluorescent microscopy (Fig-2c).

**Fig. 2:**
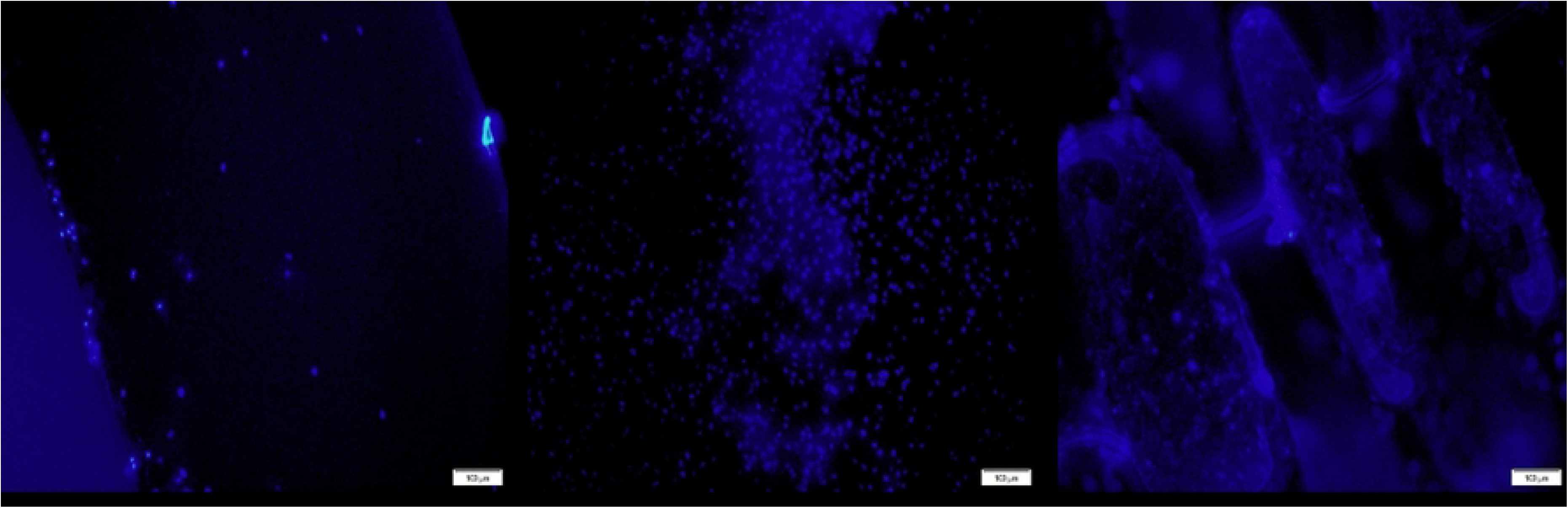
Fluorescent microscopic images of DAPI stained bacterial biofilm on (a) Empty well, (b) hydroxyapatite, (c) Titanium implant.

#### Comparison of growth of oral bacteria on TDI and HA with Empty well (Control) by real-time quantitative polymerase chain reaction

There is systematic colonization of bacterial species that include: Initial colonizers *- Streptococcus oralis* and *Actinomyces naeslundii*, Early colonizers - *Veillonella parvula*, Secondary colonizers - *Fusobacterium nucleatum* and Late colonizers - *Porphyromonas gingivalis* and *Aggregatibacter actinomycetemcomitans*. All the six bacteria were significantly more expressed on TDI than HA with p value ranging from 0.007 to 0.036 (Fig. 3)

**Fig. 3:**
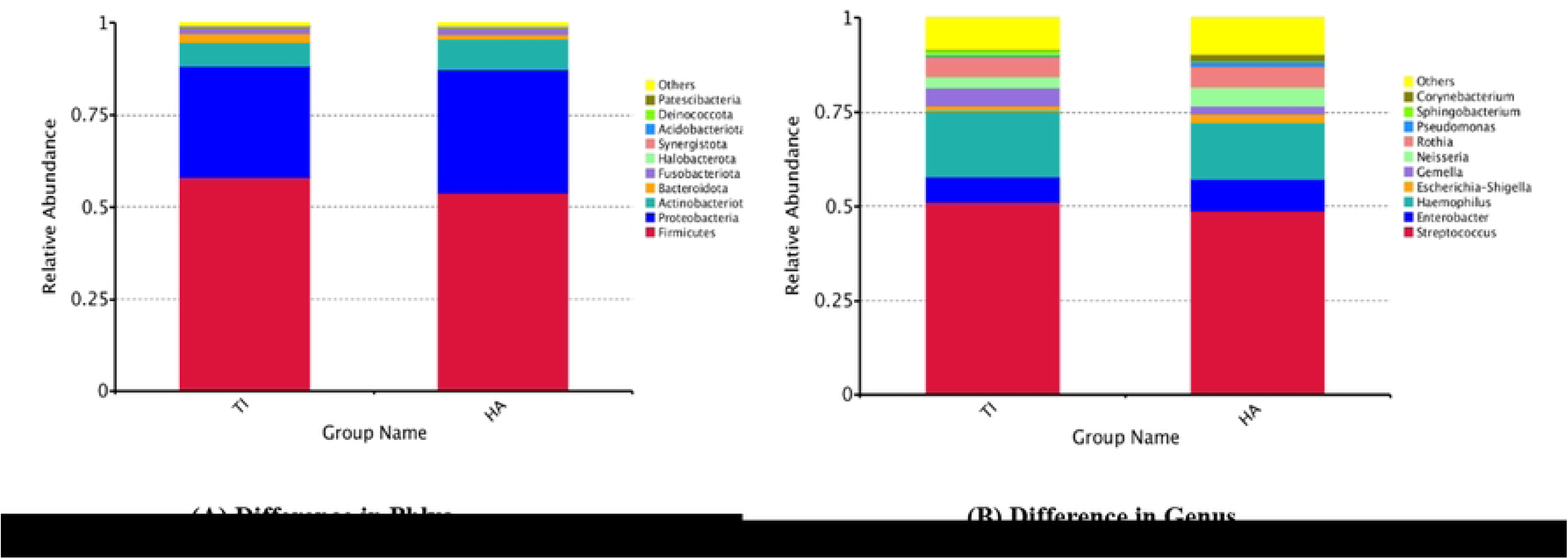
Fold expression of SO - Streptococcus oralis, AN - Actinomyces naeslundii, VP - Veillonella parvula, AA - Aggregatibacter actinomycetemcomitans, PG - Porphyromonas gingivalis, FN - Fusobacterium nucleatum on TDI and HA.

#### Targeted NGS - 16SrRNA Illumina MiseQ sequencing – Relative abundance in Phyla between TDI and HA

The phyla Firmicutes, Proteobacteria, Actinobacterioata, and Bacteroidota emerged as the predominant taxa on both titanium and hydroxyapatite substrates (Fig. 4A). However, Firmicutes and Bacteroidota exhibited greater abundance on titanium implants compared to hydroxyapatite, while Proteobacteria and Actinobacteria were relatively less abundant on titanium compared to hydroxyapatite. Commonly observed genera on both titanium implants and hydroxyapatite included Streptococcus, Enterobacter, Haemophilus, Escherichia-Shigella, Gemella, Neisseria, Rothia, Pseudomonas, Sphingobacterium, and Corynebacterium (Fig. 4B). Notably, on titanium implants, there was an elevation in Streptococcus, Haemophilus, Gemella, and Sphingobacterium genera, accompanied by a reduction in Enterobacter, Escherichia-Shigella, Neisseria, Pseudomonas, and Corynebacterium, compared to hydroxyapatite. Analysis of alpha diversity, indicative of species richness and evenness within each sample group, revealed a significant reduction in titanium implants compared to hydroxyapatite, as depicted by the Shannon index (Fig. 5A). Beta diversity, which evaluates the differences in microbial communities based on their composition, was assessed using the Weighted Unifrac distance metric to quantify dissimilarity between titanium implants and hydroxyapatite. Weighted Unifrac distance, a commonly utilized phylogenetic measurement method, highlighted the beta diversity between the two sample groups – TDI and HA (Fig. 5B).

**Fig. 4:**
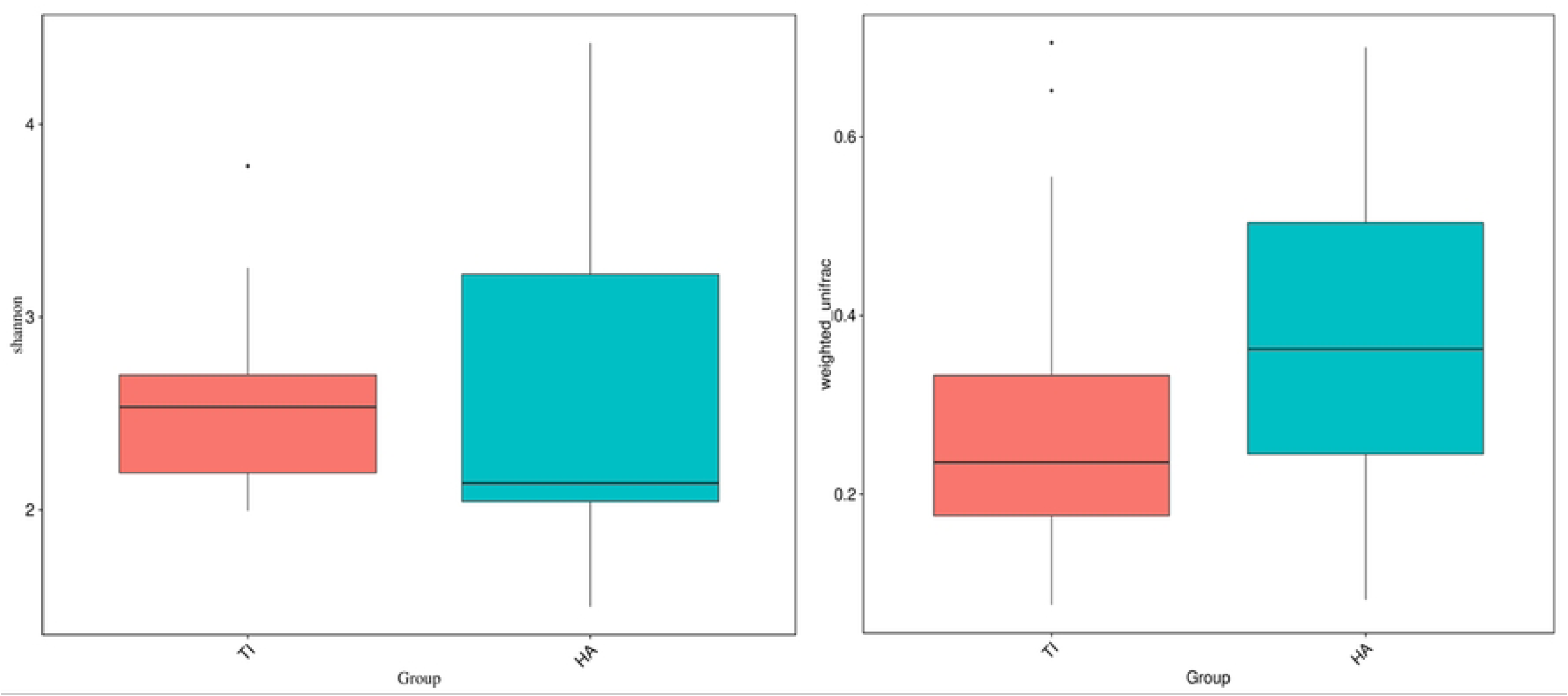
Relative abundance in Phyla and Genus between (A) TDI (Ti) and (B) HA.

**Fig. 5:**
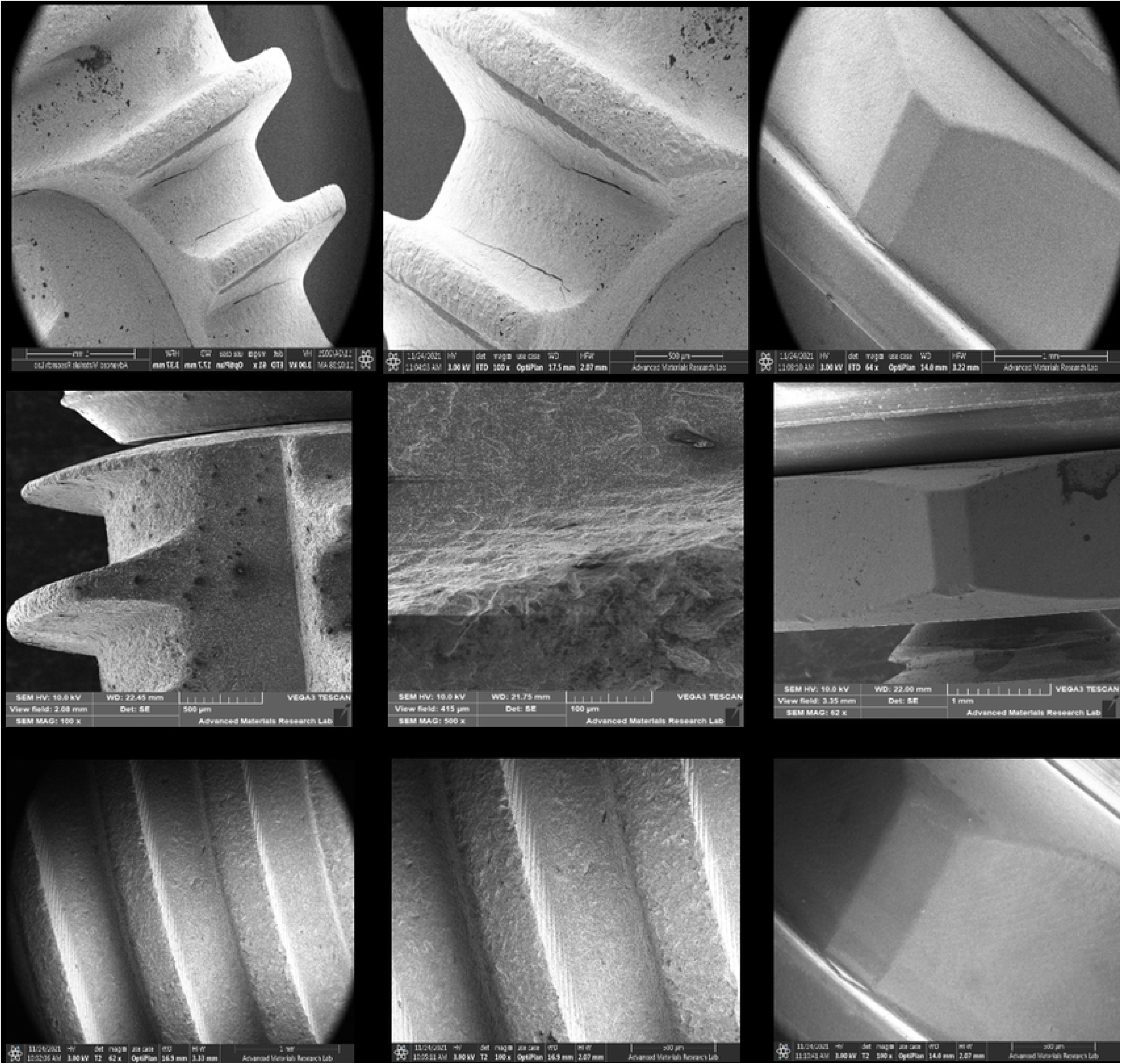
(A) Alpha diversity and (B) beta diversity between titanium implant (Ti) and hydroxyapatite (HA) indicated by the Shannon index and Weighted Unifrac distance respectively.

### In vitro studies on the effect of oral microbiota on titanium dental implants

#### Effect of bacterial polyculture of initial colonizers and late colonizers on titanium dental implant

A substantial bacterial biofilm was observed on titanium implants exposed to both sets of polyculture oral bacteria. The presence of a polyculture comprising initial colonizers, as well as one encompassing both initial and late colonizers, induced corrosion of the titanium post, deterioration of the screw thread, and damage to the abutment (Fig.6). Notably, the damage resulted was more severe in the presence of both initial colonizer, *Streptococcus mutans*, and late colonizers, namely *Porphyromonas gingivalis* and *Aggregatibacter actinomycetemcomitans*. Consequently, an increase in surface roughness enhanced the formation of thick bacterial biofilms. Furthermore, the combination of initial and late colonizers resulted in a denser biofilm compared to that formed by the sole presence of initial colonizers (Fig.6). The control titanium implant, kept immersed in Brain Heart Infusion Broth (BHIB) without bacterial inoculation, exhibited no discernible physical damage (Fig. 6).

**Fig. 6:**
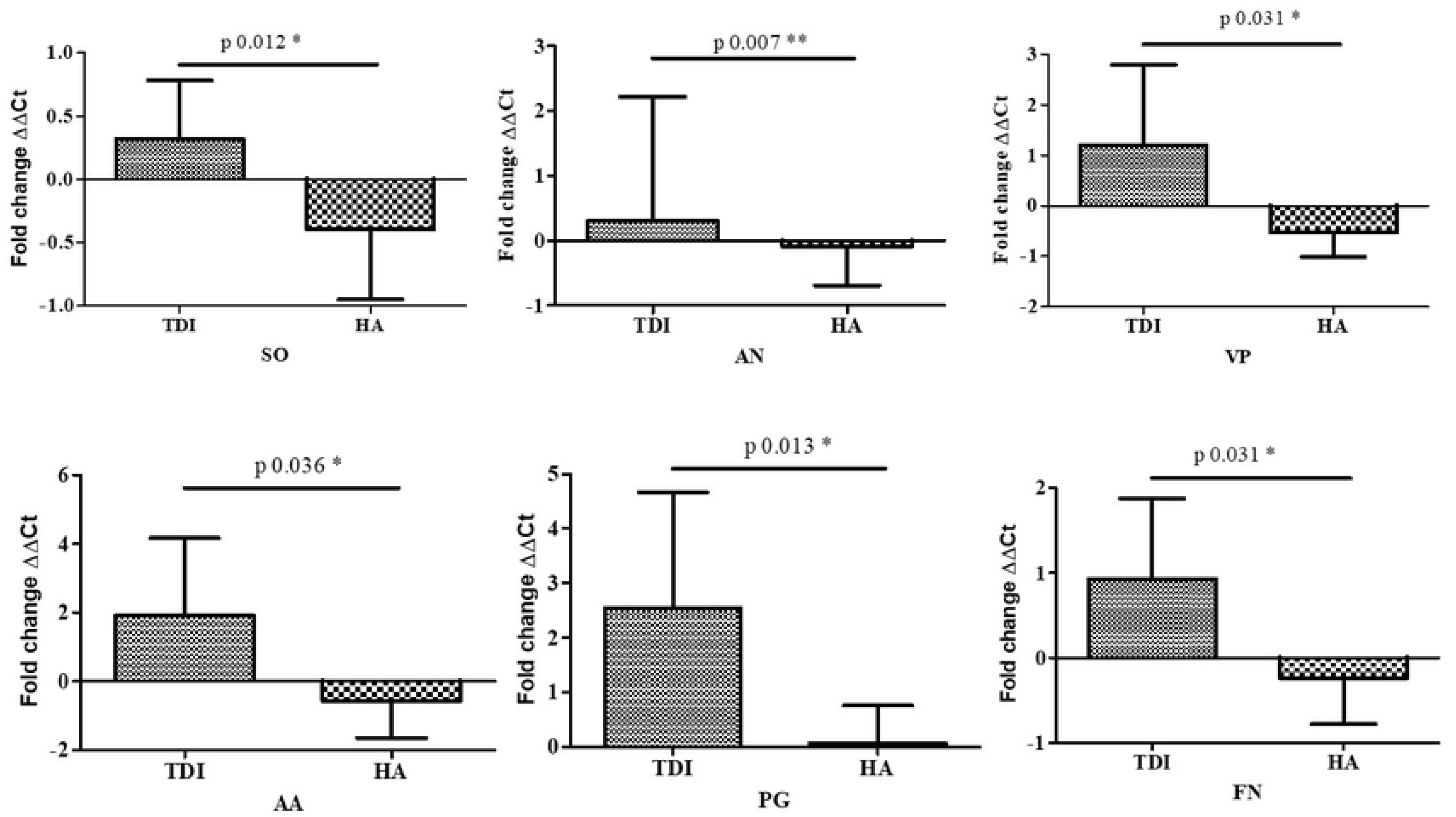
SEM of effect of set 1 - bacterial polyculture of initial colonizers, set 2 - bacterial polyculture of initial colonizers and late colonizer on titanium dental implant.

## Discussion

Dental implant-supported prostheses have been successfully restoring lost teeth for over 40 years, with a 90–95% success rate; however, failure rates of 1.9–11% have been reported [14]. These failures are attributed to factors such as microbial colonization, peri-implantitis, material corrosion, implant abutment design, and overload [15,16]. Peri-implantitis, driven by bacterial biofilm formation on implant surfaces, is the leading cause of implant-supporting tissue destruction [14,17]. This pilot study investigated the bacterial composition and its impact on titanium implant failures.

An in-house fabricated acrylic dental device with wells for hydroxyapatite, titanium implants, and empty controls enabled oral microbiota collection from gums, enamel, and implants. Participants aged 20s to 60s facilitated a diverse age range analysis. Six-hour in-vivo incubation harvested early-stage bacterial biofilms, addressing gaps in data on initial biofilm formation and non-cultivable bacteria impacting titanium implants [18,19]. This study focused on comparing the prevalence, composition, and diversity of biofilms formed during early (6h) microbial attachment on gums, teeth, and implants.

In our preliminary investigation using fluorescent microscopy and DAPI staining, biofilms were observed on plastic wells (gum control), hydroxyapatite (tooth control), and titanium implants. Biofilms on titanium implants were exceptionally dense, likely due to physicochemical properties such as surface free energy and surface roughness, with roughness being the dominant factor in bacterial adhesion [20]. Hydroxyapatite biofilms were quantifiable and denser than those in empty wells, attributed to the rapid formation of the acquired enamel pellicle. This salivary protein layer facilitates bacterial attachment to enamel surfaces despite mechanical forces [21,22,23].

Research shows that bacteria colonize the peri-implant crevice soon after implant placement, following a systematic pattern: early colonizers (e.g., *Streptococcus oralis*, *Actinomyces naeslundii*), secondary colonizers (e.g., *Veillonella parvula*), and late colonizers (e.g., *Fusobacterium nucleatum*, *Porphyromonas gingivalis*, *Aggregatibacter actinomycetemcomitans*) [24,25,26]. Our study has shown greater abundance of *Porphyromonas gingivalis* and *Aggregatibacter actinomycetemcomitans* as late colonizers within six hours of in vivo incubation, compared to early colonizers like *Streptococcus oralis* and *Veillonella parvula*, which is similar to an observation made by Siddiqui et al [27]. However, *Fusobacterium nucleatum* exhibited less growth. Unlike in vitro models, in vivo investigations incorporate host-related factors such as inflammation and pH fluctuations, offering deeper insights into implant-microbiome interactions.

An analysis of healthy oral microbiomes using 16S rDNA sequencing identified dominant phyla: Firmicutes (*Streptococcus*, *Veillonellaceae*, *Granulicatella*), Proteobacteria (*Neisseria*, *Haemophilus*), Actinobacteria (*Corynebacterium*, *Rothia*, *Actinomyces*), Bacteroidetes (*Prevotella*, *Capnocytophaga*, *Porphyromonas*), and Fusobacteria (*Fusobacterium*) [28]. Microbiota composition varies across oral niches and life stages [29]. De Melo et al found similar taxa on titanium disks, aligning with our findings from targeted NGS after 6-hour in vivo incubation on titanium implants and hydroxyapatite [30]. Firmicutes and Bacteroidota were more prevalent on titanium, with *Streptococcus* (early colonizer) and *Porphyromonas* (late colonizer) being notably present in our study. Proteobacteria and Actinobacteria were more abundant on hydroxyapatite. There was a marginal increase in the genus Streptococcus and Haemophilus on titanium implant compared to hydroxyapatite. Notably, *Gemella*, found in both healthy and periodontal conditions, was more abundant on titanium and is linked to opportunistic infections and horizontally transferrable virulence genes, possibly contributing to endocarditis [31,32,33].

Our study revealed reduced microbial diversity, richness, and evenness in the microbiome from titanium implants compared to hydroxyapatite, suggesting oral dysbiosis associated with titanium implants and potential peri-implantitis [34]. Titanium particle/ion release may contribute to this dysbiosis. Real-time qPCR and NGS analysis showed increased growth of late colonizers *Porphyromonas gingivalis* and *Aggregatibacter actinomycetemcomitans*, linked to peri-implant diseases, on titanium implants within six hours of in vivo incubation, compared to hydroxyapatite.

Gil et al. demonstrated that bacteria induce pitting corrosion on titanium surfaces under physiological in vitro conditions, degrading implant properties and potentially reducing their lifespan. However, studies on the impact of polycultures containing facultative or obligate anaerobes on titanium implants remain limited [35,36]. Our in vitro study revealed that polycultures of initial (*Streptococcus mutans*) and late colonizers (*Porphyromonas gingivalis*, *Aggregatibacter actinomycetemcomitans*) caused more severe biofilm formation and damage to titanium surfaces compared to initial colonizers alone. *S. mutans*, known for releasing lactic acid and promoting co-aggregation with periodontopathogens, facilitates corrosion, leading to surface deterioration [37,38,39]. This process releases metallic ions, exacerbating inflammation, bone resorption, and dysbiosis [40].

## Conclusion

Titanium implants, despite their clinical success, face a failure rate of 1.9% to 11%. Our in vivo and in vitro studies revealed that titanium implants can induce oral microbial dysbiosis, promoting peri-implantitis and periodontitis pathogens. Additionally, microbe-induced corrosion caused significant damage, particularly with early and late bacterial colonizers. These findings emphasize the need for further research to understand these interactions and develop strategies to enhance implant health and longevity, providing critical insights into the challenges of titanium implants in clinical practice.

